# Characterization of antiviral compounds using Bio-Layer Interferometry

**DOI:** 10.1101/2025.07.24.662752

**Authors:** Zachary C. Lorson, William M. McFadden, Grace Neilsen, Andres E. Castaner, Ryan L. Slack, Karen A. Kirby, Stefan G Sarafianos

## Abstract

Small molecule-protein interactions underpin many biological functions and play an integral role in the treatment and prevention of several human diseases. These interactions can be key to understanding the mechanism of action of these compounds. Previous methods of determining protein-protein or protein-antibody interactions have been well established; however, the use of BLI in antiviral discovery is a promising and relatively new avenue. The high-throughput nature of this method in tandem with its pM sensitivity allows for quick and seamless identification of hit compounds. Here we discuss ways to overcome common pitfalls that can occur while using BLI such as nonspecific binding (NSB) and ligand drift while offering possible solutions. Characterizing small molecule-protein interactions is not trivial and optimizing the experimental conditions is imperative. To address this gap in knowledge, we present optimized BLI protocols for the study of three cases of protein-small molecule interactions: PF74 or Lenacapavir (LEN) with HIV-1 capsid protein (CA), and Nirmatrelvir (NIR) with SARS-CoV-2 Mpro. LEN and NIR are of particular interest because they are clinically relevant, and PF74, a well-studied control, was the first compound reported to target the LEN binding site. We demonstrate that BLI can be a powerful and effective tool in calculating the binding affinities between a protein and small molecule. These newly designed methods enabled calculation of K_D_ values, the affinity between ligand and analyte, ranging from the micro to the sub-nanomolar range for CA binding events and confirmed the covalent interaction between NIR and Mpro. These protocols will facilitate efficient testing of new antivirals or derivatives in a high-throughput format.

**Summary:** Bio-Layer Interferometry (BLI) is a multifunctional technology that is used to determine valuable information on real-time kinetics including association and dissociation. Optimizing experimental conditions to acquire data about protein-ligand interactions can be challenging. We provide three example methods of collecting binding data that characterize how viral proteins interact with antivirals.

## Introduction

Discovering and developing antivirals is an expensive and laborious endeavor. Optimizing this process through efficient screening and validation methods can significantly improve output. A key aspect of drug development is determining the binding affinity of a drug candidate to its target^1, 2^. The ratio of dissociation (*k*_d_) divided by the association (*k*_a_) rate constants defines the binding constant, the affinity, expressed as K_D_. Together, *k*_a_ and *k*_d_ provide insight in how fast or slow a binding event is and how long the interaction lasts. Throughout the years, many methods have been used to determine K_D_ including microscale thermophoresis (MST)^3, 4^, isothermal titration calorimetry (ITC)^5–7^, surface plasmon resonance (SPR)^8–10^.

Bio-layer interferometry (BLI) characterizes protein and drug interactions by measuring changes in light interference patterns caused by the binding of material to the biosensor^11^. Biosensors contain two independent layers, the biocompatible layer and internal reference layer^1, 12^. The biocompatible layer contains the pre-coated immobilized molecules, such as antibodies like Anti-Penta HIS (HIS1K), metals like Ni-NTA (Ni-NTA), and other proteins like Super Streptavidin (SSA)^1^ that are used to attach the protein of interest to the biosensor. During the experiment, white light is projected down the fiber-optic sensor and is reflected from both the internal reference layer and the biocompatible layer resulting in an interference pattern. As the protein attaches to the probe, and later as the drug binds to the protein, the biocompatible layer increases in overall thickness, leading to a change of the wavelength of reflected light. This shift, affects the interference pattern of the light reflected from the two layers, altering the interference pattern recorded by the charge-coupled device (CCD) array detector, reported in nanometers (nm). This method is highly sensitive allowing for detection of ligand binding with binding affinities between 10 pM and 1 mM with high certainty,^12^ making it suitable for a wide range of antivirals to be tested with accuracy.

Other assays are used to determine binding events between two molecules. However, BLI can offer additional information or at greater ease in comparison to other methods. While MST and ITC are used to derive K_D_ values, they cannot be used to determine binding and dissociation rates, which are essential for analyzing the kinetics between the ligand and analyte. Hence, they cannot distinguish the binding differences between compounds that have similar K_D_ but drastically different *k*_a_ and *k*_d_ values (either both fast *k*_a_ and *k*_d_ rates, or both slow *k*_a_ and *k*_d_ rates resulting in the same K_D_). Unlike MST and ITC, BLI can distinguish differences in *k*_a_ and *k*_d_, which is important for the development of therapeutics that typically require long dissociation rates^2^. Unlike MST, BLI uses a label-free design, thus avoiding potential artifacts associated with attachment of additional chemical groups^13^. The combination of a label-free design, increased sensitivity, and use of lower amounts of reagents makes BLI a promising choice when aiming to characterize small molecule interactions in an enhanced throughput format^2^.

### HIV-1 Capsid

The AIDS epidemic that began in 1981 was found to be caused by a virus determined shortly thereafter as human immunodeficiency virus type 1 (HIV-1)^14^. Currently, approximately 40 million people live with HIV-1 with 1.3 million people becoming newly infected in 2023^15^.

Several iterations of drugs have been used to try and effectively neutralize the virus; however, work is still being done to find a better and long-lasting alternative to what is currently available. Problems such as access to clinics for regular drug administration, pill fatigue and overall stigma of HIV are roadblocks in treatment of this disease^14^. A novel antiviral in the treatment of HIV-1 is long-acting compound Lenacapavir (LEN)^16, 17^. This drug is extremely effective in treating drug-experienced patients due to its high potency and exceptional long-acting properties^18^. These attributes potentially offer more treatment options for patients and are an effective way of treating drug-resistant strains of this virus^19, 20^.

LEN binds to the capsid protein (CA) at the phenylalanine-glycine (FG) site, and alters the structural flexibility of the hexamer lattice, disrupting the mature CA core^21–23^. This FG-Binding domain is formed between two neighboring CA monomers at the CTD (C-terminal domain) of one monomer and the NTD (N-terminal domain) of another. The pocket plays an important role in the trafficking of capsid to the nuclear pore complex (NPC) *via* host factor interactions such as CPSF6^24^. This disruption of CA hexamer stability induced an accelerated assembly and improperly formed defective virions^23, 25^. A similar FG-targeting compound, and first reported of this class, PF-3450074 (PF74), emulates the same FG domain to bind with CA^26–28^. However, unlike LEN, PF74 is highly susceptible to degradation by cellular enzymes, but is commonly used as a benchmark or control for experiments investigating CA targeting compounds^14, 29–31^.

Experiments here with LEN and PF74 use an engineered CA that bears mutations, CA^A14C/E45C/W184A/M185A^, which enable the formation of a soluble, covalently formed hexamer through disulfide bonds (CA_HEX_)^32^. A 6x-Histidine (6xHis) fusion tag was added to the N-terminus to facilitate the hexamer attachment to the HIS1K probes; this 6xHis- tagged CA_HEX_ was expressed using *E. coli* NiCo21(DE3) and purified as reported previously^27, 33^. Additionally, buffer components were optimized to mitigate the amount of nonspecific binding (NSB) between the HIS1K probes and antivirals, adapted from^34^. While only LEN and PF74 are showcased, these two methods could be applied to various inhibitors, metabolites, and peptides.

### SARS-CoV-2 Mpro

The severe acute respiratory syndrome coronavirus 2 (SARS-CoV-2) swept across the globe in 2020 and led to the beginning of the coronavirus disease (COVID-19) pandemic^35^. As of January 2025, there have been over 700 million cases and 7 million deaths due to infection from SARS-CoV-2^36^. Thus, developing effective antivirals is of high importance.

There are several proteins critical to infection and proliferation of the virus, making them excellent targets for drug development. Among them, the main protease (Mpro) is essential for viral replication. Like other coronaviruses, SARS-CoV-2 contains a positive-sense RNA genome that can be translated after viral entry to produce two polyproteins of non-structural proteins (nsps) including those comprising the replication complex^37^. Mpro is a cysteine protease that cleaves the open reading frame 1a/b polyprotein at 11 sites resulting in the formation of the SARS-CoV-2 replication complex^36, 38, 39^. This essential activity, combined with the uniqueness of the cleavage sequence compared to human proteases, makes Mpro an exciting opportunity to target for antivirals to treat COVID-19^40^.

Three drugs have been approved by the United States FDA to fight against SARS-CoV-2. Remdesivir and Molnupiravir target the RNA-dependent RNA Polymerase, nsp12, but their use in the clinical setting have waned as more effective therapeutics became available^41, 42^. The current leading antiviral against SARS-CoV-2 is Paxlovid. Paxlovid was developed by Pfizer to target the Mpro and has proven to be extremely effective^7^. It comprises Nirmatrelvir (NIR) that targets the Mpro, as well as ritonavir, a metabolic modulator that improves the pharmacokinetic characteristics of NIR. NIR acts by forming a covalent bond with the active site cysteine of Mpro through a cyano warhead^43–45^.

Typically, validating covalent binding events, is difficult. However, using our specific probe and buffer combinations, we were able to confirm that NIR binds to Mpro covalently. Certain mutations can interrupt the covalent binding and be detected *via* BLI. For instance, the clinically relevant E166V, diminish NIR’s efficacy during SARS-CoV-2 treatment. This mutation while rare, 0.97 per million, appears in response to administration of Paxlovid and in replicon assays greatly decreases the drugs’ ability to suppress the virus^38, 46^.

These three antivirals each rely on distinct interactions with the target protein: the non-covalent PF74 and the long-acting LEN with HIV-1 capsid, and the covalent interaction of NIR and SARS-CoV-2 Mpro. Each of these examples offer a unique perspective on how antivirals interact with their targeted proteins. Here, we demonstrate the nuances of each interaction type and discuss the important experimental parameters and data analysis steps to yield consistent results in small molecule-protein studies.

### Protocol

#### Designing the Method File

##### Method 1: HIV-1 CA_HEX_-PF74

- Begin by opening a new method file and going to the first tab, Plate Definition. Here assign the wells for Equilibration, Wash, and Analyte Dissociation (columns 1,3, and 5 in Fig. 2) as Buffer by highlighting each column and designating them as Buffer. Next, assign your ligand containing wells by highlighting wells containing Protein (column 2 for rows A, B, D, F, and H in Fig. 2) and assigning them as the Load and the remainder of column 2 should be assigned as buffer.
- In column 4, rows A and B are designated as the Negative Control. Rows C, E, and G are the Reference wells and rows D, F, and H are the Sample wells.
- Moving to the Plate 1 Table, input the appropriate information into the table. For instance, the Loading wells should have the Sample ID of CA_HEX_ and the corresponding protein concentration of the hexamer, 100 µg/mL. Furthermore, both the Reference and Sample wells will need to contain the Sample ID information of the inhibitor and the concentration in µM.
- Moving to the Assay Definition tab, first create the necessary steps for the entire experiment in the Step Data List pane. First, create Baseline for 120 s and Baseline 2 for 60 s. The Loading phase for 600 s and an Association and Dissociation phase of 50 and 100 s, respectively.
- To assign wells to each step, begin by highlighting the first equilibration wells (1A/B) then double click Baseline. If done correctly, the wells will become crossed out, and the step will appear in the Assay Steps List pane. Next, highlight 2A/B and double click the Loading step. Repeat for the rest of the steps of the assay: return to 1A/B for Baseline 2, 3A/B for Baseline, 4A/B for Association, 5A/B for Dissociation.
- After completing one assay, start a new assay (if desired) by clicking the New Assay button and repeat the same steps as mentioned above for the rest of the plate (rows C/D, E/F, and G/H in Fig. 2).
- In the next tab, Sensor Assignment, highlight the wells containing sensors in the sensor tray and click Fill. This will assign which sensors will be used for the current experiment. Ensure when assigning sensors that the computer assigns the correct sensor type. If not, return to the Assay Definition tab and change the Sensor Type in the Assays Steps List pane.
- In Review Experiment, ensure that all the information inputted in previous tabs is accurate and the machine has the correct plate layout (for both trays) and run times.
- Next in the Run Experiment tab, set the experimental temperature to 25°C and the acquisition rate to Standard Kinetics. Do not change the rotations per minute of the experiment. If you wish to save the file as a method file, you do not need to provide a name or file directory. This file can then be opened, modified, and used to run future experiments.

##### Method 2: HIV-1 CA_HEX_-Lenacapavir

- Begin by opening a new method file and going to the first tab, Plate Definition. Here assign the wells for Equilibration, Wash, and Analyte Dissociation (columns 1,3, and 5 in Fig. 2) as Buffer by highlighting each column and designating them as Buffer. Next, assign your ligand containing wells by highlighting wells containing Protein (column 2 for rows A, B, D, and F in Fig. 2) and assigning them as the Load and the remainder of column 2 should be assigned as buffer.
- In column 4, rows A and B are designated as the Negative Control. Rows C and E are the Reference wells and rows D and F are the Sample wells.
- Moving to the Plate 1 Table, input the appropriate information into the table. For instance, the Loading wells should have the Sample ID of CA_HEX_ and the corresponding protein concentration of the hexamer, 200 µg/mL. Furthermore, both the Reference and Sample wells will need to contain the Sample ID information of the inhibitor and the concentration in µM.
- Moving to the Assay Definition tab, first create the necessary steps for the entire experiment in the Step Data List pane. First, create Baseline for 120 s and Baseline 2 for 60 s. The Loading phase for 300 s and an Association and Dissociation phase of 200 and 2700 s respectively.
- To assign wells to each step, begin by highlighting the first equilibration wells (1A/B) then double click Baseline. If done correctly, the wells will become crossed out, and the step will appear in the Assay Steps List pane. Next, highlight 2A/B and double click the Loading step. Repeat for the rest of the steps of the assay: return to 1A/B for Baseline 2, 3A/B for Baseline, 4A/B for Association, 5A/B for Dissociation.
- After completing one assay, start a new assay (if desired) by clicking the New Assay button and repeat the same steps as mentioned above for the rest of the plate (rows C/D and E/F in Fig. 2).
- In the next tab, Sensor Assignment, highlight the wells containing sensors in the sensor tray and click Fill. This will assign which sensors will be used for the current experiment. Ensure when assigning sensors that the computer assigns the correct sensor type. If not, return to the Assay Definition tab and change the Sensor Type in the Assays Steps List pane.
- In Review Experiment, ensure that all the information inputted in previous tabs is accurate and the machine has the correct plate layout (for both trays) and run times.
- Next in the Run Experiment tab, set the experimental temperature to 25°C and the acquisition rate to High Sensivity Kinetics. Do not change the rotations per minute of the experiment. If you wish to save the file as a method file, you do not need to provide a name or file directory. This file can then be opened, modified, and used to run future experiments.

##### Method 3: SARS-CoV-2 Mpro-NIR

- Begin by opening a new method file and going to the first tab, Plate Definition. Here assign the wells for Equilibration, Wash, and Analyte Dissociation (columns 1,3, and 5 in Fig. 2) as Buffer by highlighting each column and designating them as Buffer. Next, assign your ligand containing wells by highlighting wells containing Protein (column 2 for rows A, B, D, F and H in Fig. 2) and assigning them as the Load and the remainder of column 2 should be assigned as buffer.
- In column 4, rows A and B are designated as the Negative Control. Rows C, E, and G are the Reference wells and rows D, F, and H are the Sample wells.
- Moving to the Plate 1 Table, input the appropriate information into the table. For instance, the Loading wells should have the Sample ID of Mpro and the corresponding protein concentration of the hexamer, 100 µg/mL. Furthermore, both the Reference and Sample wells will need to contain the Sample ID information of the inhibitor and the concentration in µM.
- Moving to the Assay Definition tab, first create the necessary steps for the entire experiment in the Step Data List pane. First, create Baseline for 120 s and Baseline 2 for 60 s. The Loading phase for 400 s and an Association and Dissociation phase of 50 and 100 s respectively.
- To assign wells to each step, begin by highlighting the first equilibration wells (1A/B) then double click Baseline. If done correctly, the wells will become crossed out, and the step will appear in the Assay Steps List pane. Next, highlight 2A/B and double click the Loading step. Repeat for the rest of the steps of the assay: return to 1A/B for Baseline 2, 3A/B for Baseline, 4A/B for Association, 5A/B for Dissociation.
- After completing one assay, start a new assay (if desired) by clicking the New Assay button and repeat the same steps as mentioned above for the rest of the plate (rows C/D, E/F, G/H in Fig. 2).
- In the next tab, Sensor Assignment, highlight the wells containing sensors in the sensor tray and click Fill. This will assign which sensors will be used for the current experiment. Ensure when assigning sensors that the computer assigns the correct sensor type. If not, return to the Assay Definition tab and change the Sensor Type in the Assays Steps List pane.
- In Review Experiment, ensure that all the information inputted in previous tabs is accurate and the machine has the correct plate layout (for both trays) and run times.
- Next in the Run Experiment tab, set the experimental temperature to 25°C and the acquisition rate to Standard Kinetics. Do not change the rotations per minute of the experiment. If you wish to save the file as a method file, you do not need to provide a name or file directory. This file can then be opened, modified, and used to run future experiments.

#### Preparing Buffers and Incubating Sensors

##### Method 1: HIV-1 CA_HEX_-PF74

- Prepare a buffer of 20 mM Tris and 40 mM NaCl with additives of 20 mM Imidazole, 0.6 M Sucrose, and 1% BSA in for a total volume of 15 mL. Vortex the mixture until the BSA and Sucrose have fully dissolved into solution.
- **NOTE:** Use a tube size as close as possible to the amount of volume made to decrease the number of bubbles formed during mixing.
- Pipette 200 µL of the BLI buffer into each well that will have a sensor.
- Cover the plate with the green holder and place the sensors onto the plate using tweezers or a P10 pipette.
- **NOTE:** Ensure that the end of the sensor does not strike the plastic on exit of the original plate or upon entry of the new plate. If so, discard the sensor.
- Place the plate in the Octet machine and close the door and wait 10 minutes before continuing. After 10 minutes, begin setting up the experimental plate and have the protein on ice completely thawed.

##### Method 2: HIV-1 CA_HEX_-Lenacapavir

- Prepare a buffer of 20 mM Tris and 40 mM NaCl with additives of 20 mM Imidazole, 0.6 M Sucrose, and 1% BSA in for a total volume of 15 mL. Vortex the mixture until the BSA and Sucrose have fully dissolved into solution.
- **NOTE:** Use a tube size as close as possible to the amount of volume made to decrease the number of bubbles formed during mixing.
- Pipette 200 µL of the BLI buffer into each well that will have a sensor.
- Cover the plate with the green holder and place the sensors onto the plate using tweezers or a P10 pipette.
- **NOTE:** Ensure that the end of the sensor does not strike the plastic on exit of the original plate or upon entry of the new plate. If so, discard the sensor.
- Place the plate in the Octet machine and close the door and wait 10 minutes before continuing. After 10 minutes, begin setting up the experimental plate and have the protein on ice completely thawed.

##### Method 3: SARS-CoV-2 Mpro-NIR

- Create a buffer of 20 mM Tris pH 7.5, 100 mM NaCl, 5 mM Beta-Mercaptoethanol, 4 mM MgCl_2_ with additives of 5% Glycerol and 0.5% Tween 20 at a volume of 15 mL. Once completed, vortex the solution thoroughly until the Glycerol and Tween 20 fully dissolve.
- **NOTE:** Use a tube size as close as possible to the amount of volume made to decrease the number of bubbles formed during mixing.
- Pipette 200 µL of the BLI buffer into each well that will have a sensor, 1A-H
- Cover the plate with the green holder and place the sensors onto the plate using tweezers or a P10 pipette.
- **NOTE:** Ensure that the end of the sensor does not strike the plastic on exit of the original plate or upon entry of the new plate. If so, discard the sensor.
- Place the plate in the Octet machine and close the door and wait 15 minutes before continuing. After the 15 minutes, begin setting up the experimental plate and have the protein on ice completely thawed.

#### Setting The Experimental Plate

##### Method 1: HIV-1 CA_HEX_-PF74

- Begin by adding 198 µL of buffer into A/B 1/3-5 and 188 µL into A/B 2.
- Next adding 198 µL of buffer to C/D 1/3-5 and 198 µL to C2 and 188 µL to D2.
- Repeat this step for E/F and G/H with 199/189 µL and 199.5/189.5 µL respectively.
- Using the thawed protein aliquot, add 10 µL of protein to A/B/D/F/H ensuring the protein is mixed thoroughly.
- **NOTE:** Adding 10 µL of protein is the amount needed to get to a final protein concentration of 100 µg/mL in the 96-well plate.
- As quickly as possible, add 2 µL of DMSO into A/B 1-5 and C/D 1-3/5, 1 µL of DMSO added to E/F 1-3/5, and 0.5 µL of DMSO added to G/H 1-3/5.
- Allow drug to reach room temperature and add 2 µL to C/D 4, 1 µL to E/F 4, 0.5 µL to G/H 4. Pipette up and down multiple times and ensure the drug does not precipitate.
- **NOTE:** Adding 2/1/0.5 µL of drug resulted in a final concentration of 20/10/5 µM from a 2 mM stock.
- Place the plate into the machine and close the door. Open the method file (if needed) and ensure that the correct probes are selected in the Sensor Assignment tab. Provide the file name and directory. Finally, press GO to start data acquisition.

##### Method 2: HIV-1 CA_HEX_-Lenacapavir

- Begin by adding 198 µL of buffer into A/B 1/3-5 and 188 µL into A/B 2.
- Next adding 198 µL of buffer to C/D 1/3-5 and 198 µL to C2 and 188 µL to D2.
- Repeat this step for E/F with 199/189 µL.
- Using the thawed protein aliquot, add 10 µL of protein to A/B/D/F ensuring the protein is mixed thoroughly.
- **NOTE:** Adding 10 µL of protein is the amount needed to get to a final protein concentration of 200 µg/mL in the 96-well plate.
- As quickly as possible, add 2 µL of DMSO into A/B 1-5 and C/D 1-3/5 and 1 µL of DMSO added to E/F 1-3/5.
- Allow drug to reach room temperature and add 2 µL to C/D 4, 1 µL to E/F 4. Pipette up and down multiple times and ensure the drug does not precipitate.
- **NOTE:** Adding 2 and 1 µL of drug resulted in a final concentration of 1.25 and 0.625 µM from a 125 µM stock.
- Place the plate into the machine and close the door. Open the method file (if needed) and ensure that the correct probes are selected in the Sensor Assignment tab. Provide the file name and directory. Finally, press GO to start data acquisition.

##### Method 3: SARS-CoV-2 Mpro-NIR

- Begin by adding 199 µL of buffer into A/B 1/3-5 and 189 µL into A/B 2.
- Next adding 199 µL of buffer to C/D 1/3-5 and 199 µL to C2 and 189 µL to D2.
- Repeat this step for E/F and G/H with 199.25/189.25 µL and 199.5/189.5 µL respectively.
- Using the thawed protein aliquot, add 10 µL of protein to A/B/D/F/H ensuring the protein is mixed thoroughly.
- **NOTE:** Adding 10 µL of protein is the amount needed to get to a final protein concentration of 100 µg/mL in the 96-well plate.
- As quickly as possible, add 1 µL of DMSO into A/B 1-5 and C/D 1-3/5, 0.75 µL of DMSO added to E/F 1-3/5, and 0.5 µL of DMSO added to G/H 1-3/5.
- Allow drug to reach room temperature and add 1 µL to C/D 4, 0.75 µL to E/F 4,

0.5 µL to G/H 4. Pipette up and down multiple times and ensure the drug does not precipitate.

- **NOTE:** Adding 1/0.75/0.5 µL of drug resulted in a final concentration of 100/75/50 µM from a 20 mM stock.
- Place the plate into the machine and close the door. Open the method file (if needed) and ensure that the correct probes are selected in the Sensor Assignment tab. Provide the file name and directory. Finally, press GO to start data acquisition.

#### Analyzing the Data

##### Method 1: HIV-1 CA_HEX_-PF74

- After the run is complete, open the analyzing software and double click on the experiment file to open.
- Go to the Preprocessed Data tab and click on the Reference Sensor tab. Here you will assign the analyte only control loading wells (C2/E2/G2 in Fig. 2). Do this by selecting all three wells then right clicking and assigning them as a Reference Sensor. This action should turn the well into a diamond.
- Next, highlight a pair of experiment and analyte only controls (eg, C2/D2 in Fig. 2) and right click to Subtract Reference in Selected Sensors (select By Average in the drop down menu). This subtraction will be the first part of the double reference subtraction. Repeat for other experiment and analyte only control pairs (eg E2/F2 and G2/H2 in Fig. 2).
- Once completed, click on the Reference Sample tab. Select one of the two protein-only controls to be used and ensure that is shaped like a diamond. Next, group the diamond with the three experimental runs and again right click to Subtract Reference in Selected Sensors.
- Return to the Reference Sensor Tab and change the unused protein only control into a Reference Sensor.
- In the Data Correction tab, align the Y-axis to the Average of the Baseline Step. Furthermore, the Inter-Step Correction should be set to Baseline Step at Time 0 s. Lastly, turn on the Savitzky-Golay Filtering.

##### Method 2: HIV-1 CA_HEX_-Lenacapavir

- After the run is complete, open the analyzing software and double click on the experiment file.
- Go to the Preprocessed Data tab and click on the Reference Sensor tab. Here you will assign the drug only controls, C2 and E2. Do this by selecting all three wells then right clicking and assigning them as a Reference Sensor. This action should turn the well into a diamond.
- Next, highlight the experimental and drug only control in pairs, C/D, and right click to Subtract Reference in Selected Sensors. This subtraction will be the first part of the double reference subtraction.
- Once completed, click on the Reference Sample tab. Select one of the two protein-only controls to be used and ensure that is shaped like a diamond. Next, group the diamond with the three experimental runs and again right click to Subtract Reference in Selected Sensors.
- Return to the Reference Sensor Tab and change the unused protein only control into a Reference Sensor.
- In the Data Correction tab, align the Y-axis to the Average of the Baseline Step. Furthermore, the Inter-Step Correction should be set to Baseline Step at Time 0 s. Lastly, turn on the Savitzky-Golay Filtering.

##### Method 3: SARS-CoV-2 Mpro-NIR

- After the run is complete, open the analyzing software and double click on the experiment file.
- Go to the Preprocessed Data tab and click on the Reference Sensor tab. Here you will assign the drug only controls, C2/E2/G2. Do this by selecting all three wells then right clicking and assigning them as a Reference Sensor. This action should turn the well into a diamond.
- Next, highlight the experimental and drug only control in pairs, C/D, and right click to Subtract Reference in Selected Sensors. This subtraction will be the first part of the double reference subtraction.
- Once completed, click on the Reference Sample tab. Select one of the two protein-only controls to be used and ensure that is shaped like a diamond. Next, group the diamond with the three experimental runs and again right click to Subtract Reference in Selected Sensors.
- Return to the Reference Sensor Tab and change the unused protein only control into a Reference Sensor.
- In the Data Correction tab, align the Y-axis to the Average of the Baseline Step. Furthermore, the Inter-Step Correction should be set to Baseline Step at Time 0 s. Lastly, turn on the Savitzky-Golay Filtering.

## REPRESENTATIVE RESULTS

Generally, there are five fundamental steps: 1) Baseline, 2) Loading, 3) Baseline 2, 4) Association, 5) Dissociation (Figure 1). Figure 2 displays a general plate layout of a typical BLI experiment to achieve this. The initial Baseline occurs before the ligand binds the sensor and is represented as light blue (Figure 2). This step allows the probes time to equilibrate before protein is introduced. Usually, between 120 to 180 s is sufficient time for the probe-buffer interaction to become stable and plateau prior to continuing with the experiment.

**Figure 1.**
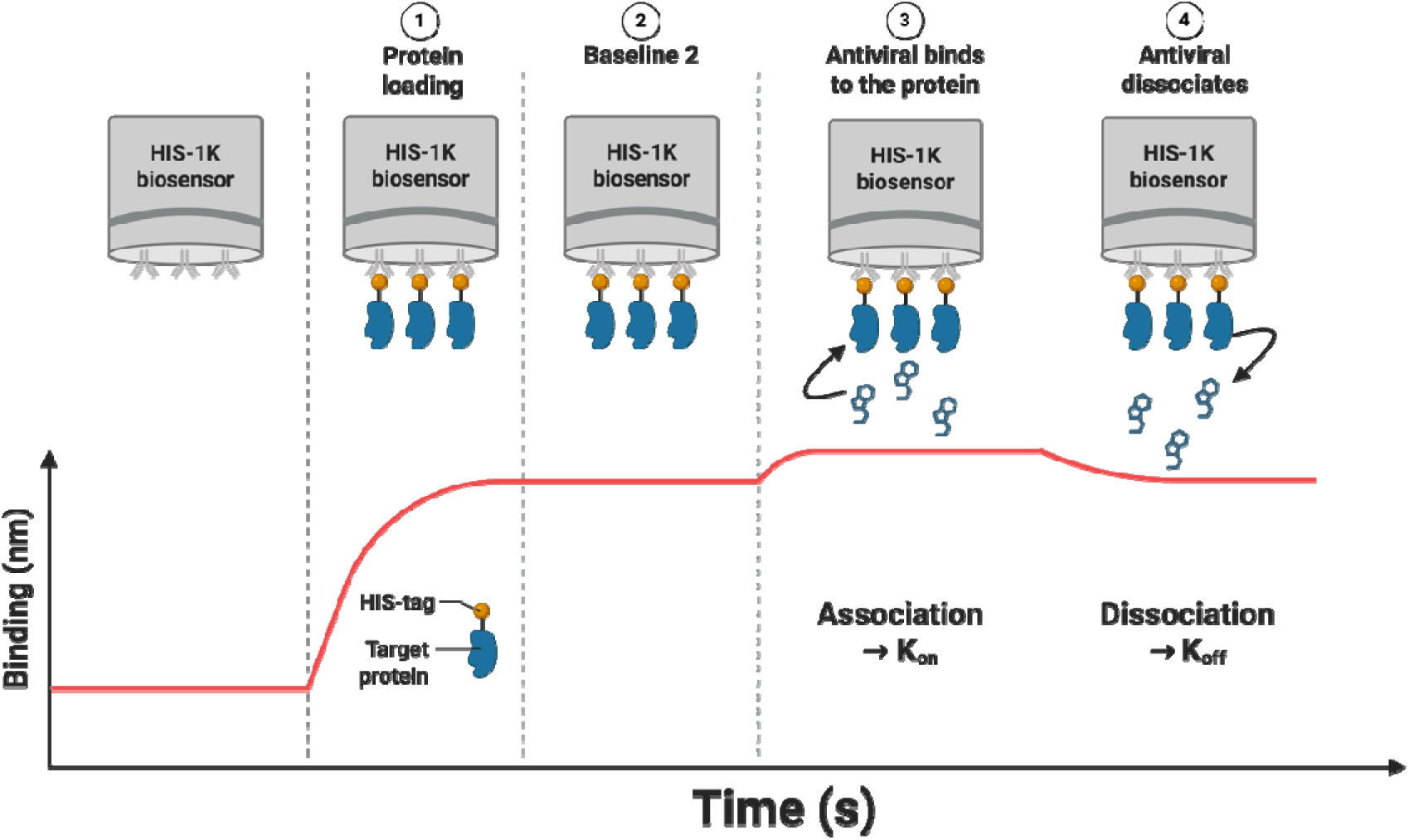
Cartoon representation of a typical BLI experiment. Hydrated probes are first equilibrated in buffer at baseline. Then in the loading phase, protein (CA_HEX_ or Mpro) binds to the sensor, followed by a short baseline in buffer. In association, the antiviral of choice (PF74 or LEN for CA_HEX_, NIR for Mpro) binds to the protein and loses interaction in the dissociation phase. Made in BioRender.

**Figure 2.**
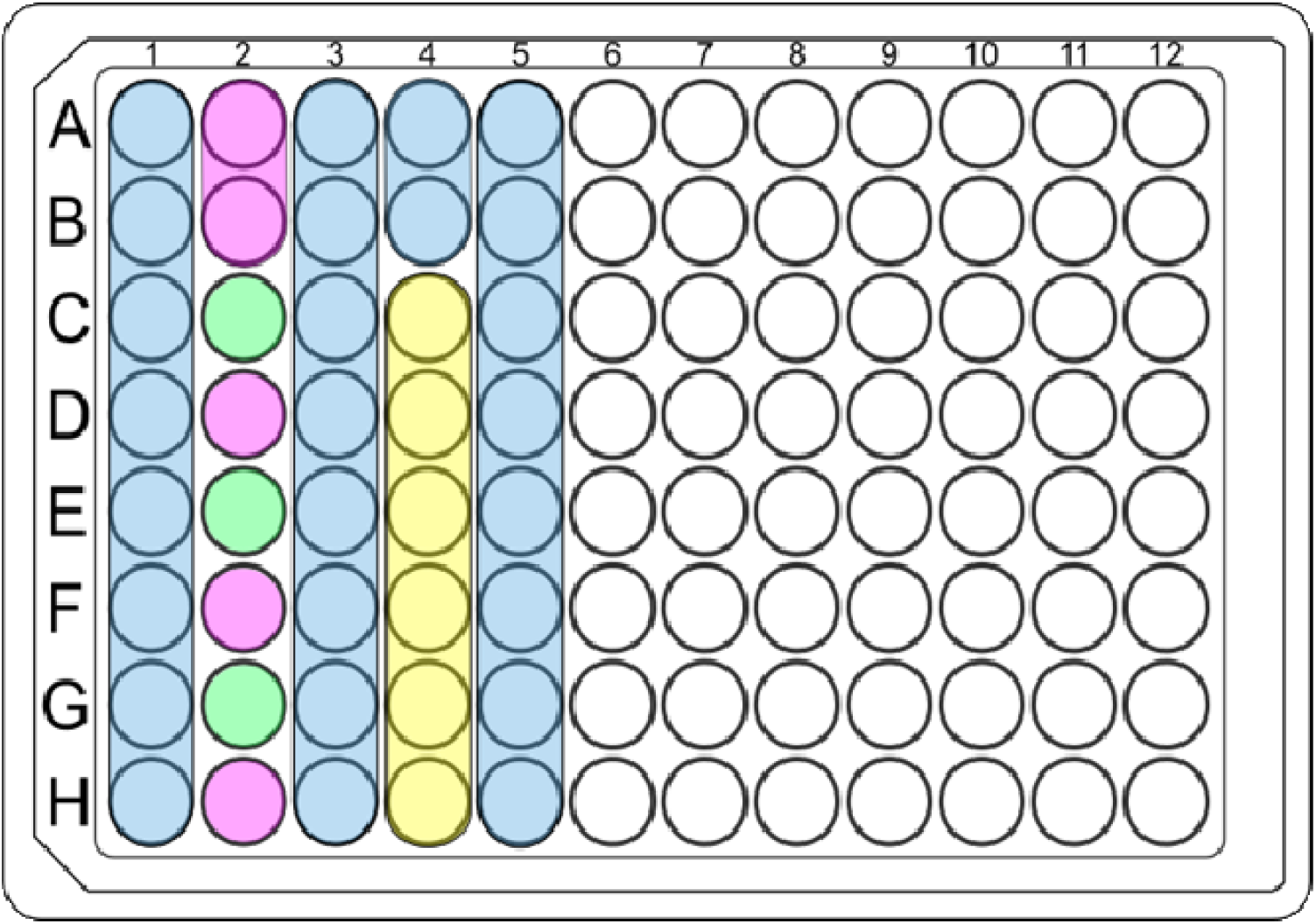
Schematic of 96-well sample plate. Plate layout of a procedure using the Sartorius Octet® R2 system. The buffer only wells (light blue) are used for pre-loading equilibration, post-loading equilibration, and dissociation. Protein loading wells (light pink) contain 100 µg/mL and protein absent wells (light green) are the analyte only controls. Rows A and B function as the protein only controls with no analyte being present, and rows D, F, and H are the experimental runs. The analyte-containing wells (light yellow) contain descending concentrations of antiviral.

The Loading step occurs in column 2 and is where the protein is immobilized to the specific sensor type, Ni-NTA or HIS1K. Each protein and probe interaction is unique in how much protein is loaded and how long it takes to reach saturation^12^. However, a protein concentration of 50-100 µg/mL is a good starting point with a time of 300 to 600 s, if binding is not known. These concentrations yielded a consistent amount of protein being loaded onto the probe in a short amount of time. However, lower concentrations, not shown, did not fully saturate the probe and increased drift in the experiment. For CA_HEX_ and HIS1K experiments, there should be between 1.4-1.6 nm worth of protein loaded onto the sensor. Shifts less than 1nm, should be considered unreliable due to a faulty probe or misfolded protein. For Mpro and Ni-NTA experiments, there should be a shift between 6-8 nm with loading under 4 nm being considered unreliable as well. Inconsistent loading of the protein in the initial steps can lead to inaccuracy in the downstream steps.

For all experiments shown, protein drift should be greatly reduced with a change of +/- 0.04 nm during the second baseline step. Any significant drift seen in this step could be indicative of a buffer mismatch, probe malfunction or misfolded protein. In this context, protein drift can be defined as a gradual, but significant, increase or decrease in signal while nothing is being added or taken away from the probe.

For the PF74 studies, three concentrations were used, 20, 10, and 5 µM with an average shift of 0.04 nm after performing subtraction (Figure 3). Using the long-acting LEN required lower antiviral concentrations at 1.25, 0.625, and 0.3125 µM to minimize background and to be able to detect dissociation. A shift of 0.03-0.05 nm should occur within 100 s and then remain flat (Figure 4). Association of NIR on average resulted in a shift of approximately 0.06-0.08 nm and plateauing within 50 s, with antiviral concentrations from 100, 75, 50 µM (Figure 5).

**Figure 3.**
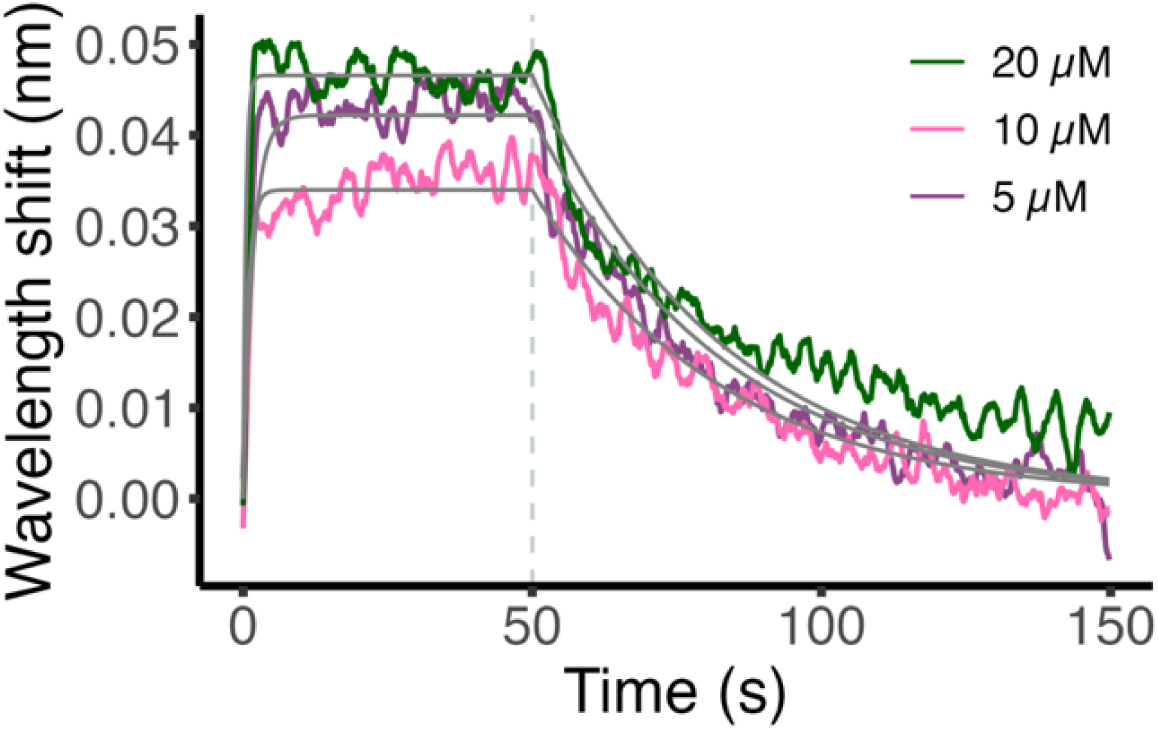
Stacked graph of analyzed data from CA_HEX_:PF74 experiment. Data were processed as described in the text and fit with a 1:1 model. Three representative curves are shown of the concentrations used, 20, 10, 5 µM. Global fitting results of the nine runs: *K*_D_ = 365 ± 0.17 nM, *k*_a_ = 9.68 x 10^4^ ± 0.43 M^-1^ s^-1^, *k*_d_ = 3.53 x 10^-2^ ± 0.03 s^-1^. Goodness of fit R^2^ = 0.881.

**Figure 4.**
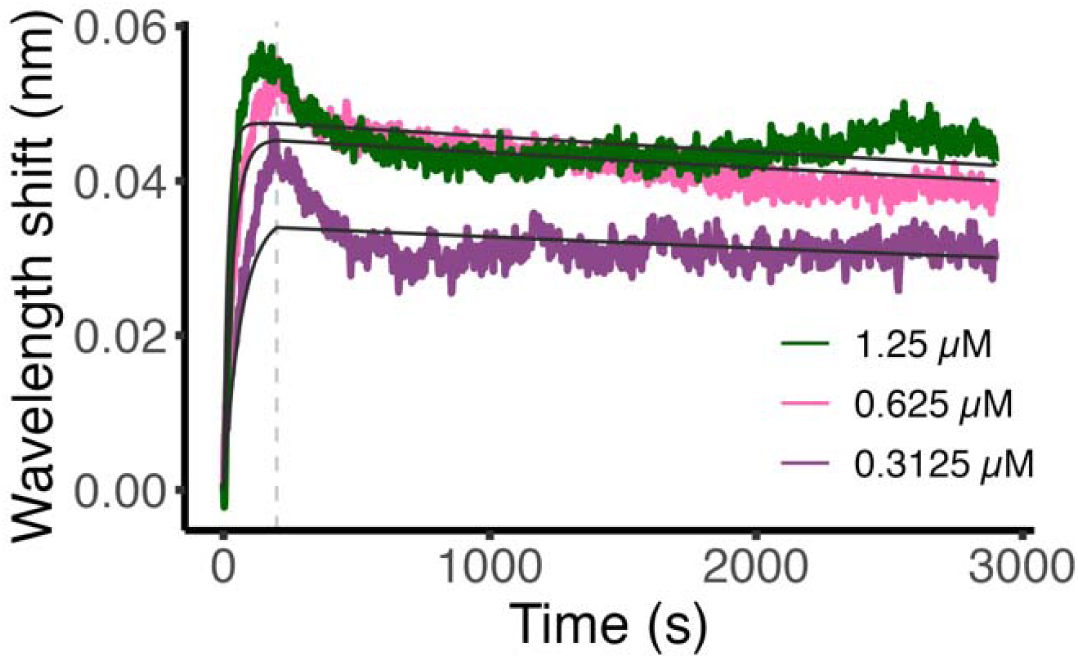
Stacked graph of analyzed data from CA_HEX_:LEN experiment. Data were processed as described in the text and fit with a 1:1 model. Three representative curves are shown of the concentrations used, 1.25, 0.625, 0.3125 µM. Global fitting results of the nine runs: *K*_D_ = 0.89 ± 0.15 nM, *k*_a_ = 4.93 x 10^4^ ± 0.05 M^-1^ s^-1^, *k*_d_ = 4.41 x 10^-5^ ± 0.06 s^-1^. Goodness of fit R^2^ = 0.845. As similarly reported in Bruce *et al.*, (2024)^27^.

**Figure 5.**
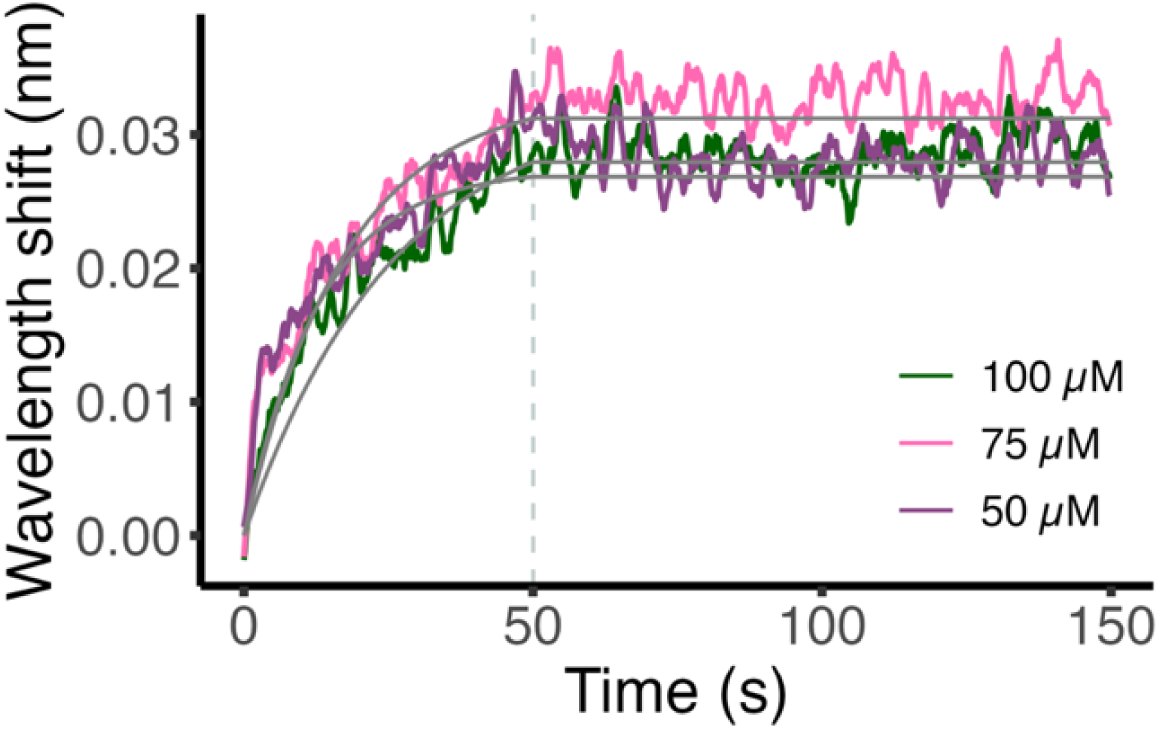
Stacked graph of analyzed data from Mpro:NIR experiment. Data were processed as described in the text and fit with a 1:1 model. Three representative curves are shown of the covalent interaction with concentrations of, 100, 75, 50 µM. Global fitting results of the nine runs: *K*_D_ = 1.26 nM, *k*_a_ = 7.74 x 10^2^ ± 0.09 M^-1^ s^-1^, *k*_d_ = 9.77 x 10^-7^ s^-1^. Goodness of fit R^2^ = 0.873. As reported in Neilsen *et al.*, (2025)^38^.

For antivirals that have faster dissociation rates, such as PF74, it is best to use typical dissociation times of 100 s to capture the full dissociation event. However, for antivirals such as LEN, the dissociation step was extended to 2700 s to allow for proper calculation of dissociation. During the dissociation period for Mpro, there should be little to no loss of interaction between the drug and protein after 100 s after subtracting both controls. The dissociation phase for NIR was shorter in comparison to LEN because of the drift issues experienced while using the Ni-NTA probes. Thus, to make the experiment more reproducible, the dissociation rate was set to 100 s.

To accurately model the antiviral binding to the protein of interest, both the drift of the protein and the background signal of the antiviral to the probe must be accounted for. We attempted to account for these issues by using double reference subtraction. For protein drift, we used two protein-only controls for each experiment while only choosing one control to be subtracted after the end of the run. This subsequently led us to predominately use the same probe as our experimental runs, the one with similar drift patterns. Furthermore, we used an antiviral control that was run at the same time and concentration as the equivalent experimental run. In all runs, both of these controls were subtracted from the experimental groups.

To better analyze the data, we used two alignment shifts: Aligning to the Y-axis, and an inter-step correction. Aligning the Y-axis to the baseline step creates the same start point before starting the association step, allowing for easier interpretation of the association phase. Using an inter-step correction to Baseline helps account for any buffer mismatching that occurs between the association and dissociation phases. To be able to use the global fit function, we used a series of antiviral concentrations, yielding more accurate kinetic constants. Furthermore, determining the overall quality of the data can be difficult when looking at new interactions between proteins and antivirals. While concentrations used are significantly higher than the *K*_D_ reported, this was done to attain a reliable response across all experiments.

## Discussion

BLI offers a user-friendly experience in developing and executing methods to study *in vitro* biomolecular interactions. The ease in operation of the machine and the relative low cost compared to SPR offers a competitive advantage in targeting small molecule interactions with proteins. While there have been reports that characterize the kinetics of interactions for both CA_HEX_ and Mpro, most experiments have either utilized SPR or have focused on interactions outside of protein-small molecules^2, 9, 23, 25, 47^. This leaves a significant moat of knowledge to be addressed using this new technology. More specifically, the methods provided here can quickly and effectively be used to test binding of compounds to HIV-1 CA and SARS-CoV-2 Mpro. Using these methods, we characterized the binding affinities of three compounds, each having a distinct binding profile: covalent binding, or non-covalent with either low or high affinity. These binding modalities cover most types of small molecule interactions and enabling efficient analysis. Lastly, optimizing the buffer conditions and creating a reliable method can be a time-consuming part of creating a BLI experiment. By troubleshooting these issues for two clinically relevant proteins, screening a set of antivirals against these proteins should be easier for future studies utilizing BLI systems.

Due to the components in the buffers, primarily the BSA, sucrose, and glycerol, it is critical to make the BLI running buffer fresh each day. If an old buffer is used, the chance of protein instability and NSB increases. Decreasing the amount of noise and buffer differences is imperative for sensitive measurements of small molecule interactions which can be greatly affected by DMSO. Furthermore, while setting the tray it is important to add DMSO to each well individually. Another crucial step while designing the method, is matching which sensor is being used during the experiment and using the same for the protein-only control. There was a consistent divergence in how the probes behaved in long-lasting experiments. This was especially noticeable in the protein-only controls. There was upward drift in the top probe, whereas the bottom probe stayed steady for the entire dissociation phase. For this reason, it is recommended that the protein-only control and experimental runs are performed using the same probe to minimize overall error. While this issue might be on a case-by-case basis, it is important to remain vigilant when analyzing the data.

From our experience, a limitation of testing small molecule inhibitors against proteins is the low margin of error for each component of the experiment. Each experimental component, the protein-only and drug-only controls and the experimental run, can have an unusual drift or buffer jump that makes the data unusable. This is more of an issue for small molecules in comparison to protein-protein or protein-antibody interactions because of the size of the nm shift when association occurs. For small molecules, a shift between 0.04 - 0.10 nm is typical for these settings. However, protein-protein interactions usually have a significantly larger nm shift making buffer jumps or drift less impactful to the overall data. For example, the Mpro experiments require the protein-only control to have little to no drift during the dissociation period to generate valid results. However, there were multiple instances where these controls would fail to stay flatlined and would deviate significantly in the positive or negative direction. We found that the chance of drift occurring increases in longer experiments, likely due to dehydration. Thus, it is suggested that experiments are restricted to approximately 100 seconds of dissociation. However, in extreme cases of sub-nanomolar affinity, longer dissociation times are suggested for more accurate assessment of binding constants.

Different buffers may be tested as the buffer type can greatly influence the amount of protein loaded onto the sensor, binding stability, and NSB. Additives as well as changes to the pH can also be tested towards addressing potential issues. The running buffer conditions were fine-tuned were for the SARS-CoV-2 Mpro experiments. In these experiments, we used a previously published buffer^34^, but noted a non-linear decay of the protein-sensor interaction that made subtraction of the protein-only control difficult. Hence, we used a series of buffers containing different detergents and other additives widely known to help stabilize proteins and reduce overall NSB interactions: Tween 20, glycerol, BSA, imidazole, and sucrose. Using varying concentrations and combinations of these additives, seven runs were performed. The optimal conditions contained 0.5% Tween 20 and 5% glycerol and substantially decreased the degradation of the protein-probe interaction. Additionally, it was equal to or better than all other conditions tested in suppressing NSB. By reducing the protein loss to less than 0.1 nm it allowed for a much more reliable protein-only control subtraction.

Another potential issue that may arise is the liquid evaporation in wells during experiments over 100 seconds of dissociation. Notably, the R2 system does not have a lid to cover the experimental plate that decreases evaporation and may not be as effective for experiments that last more than three hours. Efforts to increase the ambient humidity, resulted in more consistent results measurements.

While these issues and limitations can be initially troublesome, BLI can still be an effective tool in calculating the binding affinities between proteins and small molecules. Utilizing different buffer conditions, sensor types, and method design we were able to characterize three distinct small molecule-protein interactions. These methods are useful in the development and improvement of antivirals and can be applied to other small molecules and proteins.

## Acknowledgments

This research was supported in part by the National Institutes of Health (R01 AI120860, R01 AI167356, P30 AI050409, and U54 AI170855 to S.G.S.; R21 AI189247 to K.A.K.; F31 AI179181 to G.N.; F31 AI174951 to W.M.M.; G.N. and W.M.M. were supported in part by T32 GM135060; G.N. was supported in part by T32 AI106699; R.L.S. supported in part by T32 AI157855). The content is solely the responsibility of the authors and does not necessarily represent the official views of the National Institutes of Health. W.M.M. thanks ARCS Atlanta for their support. S.G.S. acknowledges funding from the Nahmias-Schinazi Distinguished Chair in Research. The plasmid encoding the untagged Disulfide Stabilized CA Hexamers (CA_HEX_) was provided by Dr. Owen Pornillos (University of Utah).

## Author contributions

Conceptualization, Z.C.L., W.M.M., G.N., and S.G.S.; methodology & investigation, Z.C.L., W.M.M., G.N., and R.L.S.; visualization, Z.C.L., G.N., and W.M.M.; resources, Z.C.L., A.E.C., and K.A.K.; writing – original draft, Z.C.L., G.N., and W.M.M.; writing – review & editing Z.C.L., W.M.M., G.N., R.L.S., K.A.K., and S.G.S.; funding acquisition, K.A.K., and S.G.S..

## References

1. Murali, S., Rustandi, R.R., Zheng, X., Payne, A., Shang, L. Applications of Surface Plasmon Resonance and Biolayer Interferometry for Virus–Ligand Binding. Viruses. 14 (4), 717, doi: 10.3390/v14040717 (2022).

2. Jug, A., Bratkovič, T., Ilaš, J. Biolayer interferometry and its applications in drug discovery and development. TrAC Trends in Analytical Chemistry. 176, 117741, doi: 10.1016/j.trac.2024.117741 (2024).

3. El Deeb, S., et al. Microscale thermophoresis as a powerful growing analytical technique for the investigation of biomolecular interaction and the determination of binding parameters. Methods and Applications in Fluorescence. 10 (4), 042001, doi: 10.1088/2050-6120/ac82a6 (2022).

4. Nasreddine, R., Nehmé, R. Microscale thermophoresis for studying protein-small molecule affinity: Application to hyaluronidase. Microchemical Journal. 170, 106763, doi: 10.1016/j.microc.2021.106763 (2021).

5. Di Trani, J.M. et al. Rapid measurement of inhibitor binding kinetics by isothermal titration calorimetry. Nature Communications. 9 (1), 893, doi: 10.1038/s41467-018-03263-3 (2018).

6. Su, H., Xu, Y. Application of ITC-Based Characterization of Thermodynamic and Kinetic Association of Ligands With Proteins in Drug Design. Frontiers in Pharmacology. 9, 1133, doi: 10.3389/fphar.2018.01133 (2018).

7. Perozzo, R., Folkers, G., Scapozza, L. Thermodynamics of Protein–Ligand Interactions: History, Presence, and Future Aspects. Journal of Receptors and Signal Transduction. 24 (1–2), 1–52, doi: 10.1081/RRS-120037896 (2004).

8. Miyazaki, C.M., Shimizu, F.M., Mejía-Salazar, J.R., Oliveira, O.N., Ferreira, M. Surface plasmon resonance biosensor for enzymatic detection of small analytes. Nanotechnology. 28 (14), 145501, doi: 10.1088/1361-6528/aa6284 (2017).

9. Li, H. et al. Inhibitory Effects and Surface Plasmon Resonance Based Binding Affinities of Dietary Hydrolyzable Tannins and Their Gut Microbial Metabolites on SARS-CoV-2 Main Protease. Journal of agricultural and food chemistry. 69 (41), 12197–12208, doi: 10.1021/acs.jafc.1c03521 (2021).

10. Phillips, K.S., Cheng, Q. Recent advances in surface plasmon resonance based techniques for bioanalysis. Analytical and Bioanalytical Chemistry. 387 (5), 1831– 1840, doi: 10.1007/s00216-006-1052-7 (2007).

11. Concepcion, J. et al. Label-free detection of biomolecular interactions using BioLayer interferometry for kinetic characterization. Combinatorial Chemistry & High Throughput Screening. 12 (8), 791–800, doi: 10.2174/138620709789104915 (2009).

12. Apiyo, D. Biomolecular Binding Kinetics Assays on the Octet® BLI Platform. at <https://www.sartorius.com/resource/blob/742330/05671fe2de45d16bd72b8078ac28980d/octet-biomolecular-binding-kinetics-application-note-4014-en-1--data.pdf> (2022).

13. Petersen, R.L. Strategies Using Bio-Layer Interferometry Biosensor Technology for Vaccine Research and Development. Biosensors. 7 (4), 49, doi: 10.3390/bios7040049 (2017).

14. M. Ravichandran, S., M. McFadden, W., A. Snyder, A., G. Sarafianos, S. State of the ART (antiretroviral therapy): Long-acting HIV-1 therapeutics. Global Health & Medicine. 6 (5), 285–294, doi: 10.35772/ghm.2024.01049 (2024).

15. The path that ends AIDS: UNAIDS Global AIDS Update 2023. Geneva: Joint United Nations Programme on HIV/AIDS; 2023. Licence: CC BY-NC-SA 3.0 IGO.

16. Kelley, C.F. et al. Twice-Yearly Lenacapavir for HIV Prevention in Men and Gender-Diverse Persons. New England Journal of Medicine. 392 (13), 1261–1276, doi: 10.1056/NEJMoa2411858 (2025).

17. McFadden, W.M., Faerch, M., Kirby, K.A., Dick, R.A., Torbett, B.E., Sarafianos, S.G. Considerations for capsid-targeting antiretrovirals in pre-exposure prophylaxis. Trends in Molecular Medicine. (25), 1–13, doi: 10.1016/j.molmed.2025.01.013 (2025).

18. Ogbuagu, O. et al. Efficacy and safety of the novel capsid inhibitor lenacapavir to treat multidrug-resistant HIV: week 52 results of a phase 2/3 trial. The Lancet HIV. 10 (8), e497–e505, doi: 10.1016/S2352-3018(23)00113-3 (2023).

19. Pinto, M.F., Sirina, J., Holliday, N.D., McWhirter, C.L. High-throughput kinetics in drug discovery. SLAS Discovery. 29 (5), 100170, doi: 10.1016/j.slasd.2024.100170 (2024).

20. Sohraby, F., Nunes-Alves, A. Advances in computational methods for ligand binding kinetics. Trends in Biochemical Sciences. 48 (5), 437–449, doi: 10.1016/j.tibs.2022.11.003 (2023).

21. Bester, S.M. et al. Structural and mechanistic bases for a potent HIV-1 capsid inhibitor. *Science (New York*, N.Y.). 370 (6514), 360–364, doi: 10.1126/science.abb4808 (2020).

22. McFadden, W.M. et al. Rotten to the core: antivirals targeting the HIV-1 capsid core. Retrovirology. 18 (1), 41, doi: 10.1186/s12977-021-00583-z (2021).

23. Link, J.O. et al. Clinical targeting of HIV capsid protein with a long-acting small molecule. Nature. 584 (7822), 614–618, doi: 10.1038/s41586-020-2443-1 (2020).

24. Zhuang, S., Torbett, B.E. Interactions of HIV-1 Capsid with Host Factors and Their Implications for Developing Novel Therapeutics. Viruses. 13 (3), 417, doi: 10.3390/v13030417 (2021).

25. Bester, S.M. et al. Structural and Mechanistic Bases of Viral Resistance to HIV-1 Capsid Inhibitor Lenacapavir. mBio. 13 (5), e01804–22, doi: 10.1128/mbio.01804-22.

26. Bhattacharya, A. et al. Structural basis of HIV-1 capsid recognition by PF74 and CPSF6. Proceedings of the National Academy of Sciences of the United States of America. 111 (52), 18625–18630, doi: 10.1073/pnas.1419945112 (2014).

27. Bruce, A. et al. A Tag-Free Platform for Synthesis and Screening of Cyclic Peptide Libraries. Angewandte Chemie (International Ed. in English). 63 (21), e202320045, doi: 10.1002/anie.202320045 (2024).

28. Gres, A.T., Kirby, K.A., KewalRamani, V.N., Tanner, J.J., Pornillos, O., Sarafianos, S.G. X-ray crystal structures of native HIV-1 capsid protein reveal conformational variability. Science. 349 (6243), 99–103, doi: 10.1126/science.aaa5936 (2015).

29. Vernekar, S.K.V. et al. Toward Structurally Novel and Metabolically Stable HIV-1 Capsid-Targeting Small Molecules. Viruses. 12 (4), 452, doi: 10.3390/v12040452 (2020).

30. Xu, S., Sun, L., Ding, D., Zhang, X., Liu, X., Zhan, P. Metabolite Identification of HIV- 1 Capsid Modulators PF74 and 11L in Human Liver Microsomes. Metabolites. 12 (8), 752, doi: 10.3390/metabo12080752 (2022).

31. Price, A.J. et al. Host Cofactors and Pharmacologic Ligands Share an Essential Interface in HIV-1 Capsid That Is Lost upon Disassembly. PLOS Pathogens. 10 (10), e1004459, doi: 10.1371/journal.ppat.1004459 (2014).

32. Pornillos, O. et al. X-ray Structures of the Hexameric Building Block of the HIV Capsid. Cell. 137 (7), 1282–1292, doi: 10.1016/j.cell.2009.04.063 (2009).

33. McFadden, W.M. et al. Modifying PF74 Improves Anti-HIV-1 Activity Against the Resistance-Associated Capsid Mutation N74D. [CROI Abstract 727]. Topics in Antiviral Medicine. 33 (1), 208 (2025).

34. Dubrow, A., Zuniga, B., Topo, E., Cho, J.-H. Suppressing Nonspecific Binding in Biolayer Interferometry Experiments for Weak Ligand–Analyte Interactions. ACS Omega. 7 (11), 9206–9211, doi: 10.1021/acsomega.1c05659 (2022).

35. Lotfi, M., Rezaei, N. SARS-CoV-2: A comprehensive review from pathogenicity of the virus to clinical consequences. Journal of Medical Virology. 92 (10), 1864–1874, doi: 10.1002/jmv.26123 (2020).

36. Neilsen, G. et al. Dimming the corona: studying SARS-coronavirus-2 at reduced biocontainment level using replicons and virus-like particles. mBio. 15 (12), e03368–23, doi: 10.1128/mbio.03368-23 (2024).

37. V’kovski, P., Kratzel, A., Steiner, S., Stalder, H., Thiel, V. Coronavirus biology and replication: implications for SARS-CoV-2. Nature Reviews Microbiology. 19 (3), 155– 170, doi: 10.1038/s41579-020-00468-6 (2021).

38. Neilsen, G. et al. Strategy to overcome a nirmatrelvir resistance mechanism in the SARS-CoV-2 nsp5 protease. Science Advances. 11 (23), eadv8875, doi: 10.1126/sciadv.adv8875 (2025).

39. Zheng, Y. et al. SARS-CoV-2 NSP5 and N protein counteract the RIG-I signaling pathway by suppressing the formation of stress granules. Signal Transduction and Targeted Therapy. 7 (1), 1–12, doi: 10.1038/s41392-022-00878-3 (2022).

40. Zhang, L. et al. Crystal structure of SARS-CoV-2 main protease provides a basis for design of improved α-ketoamide inhibitors. Science. 368 (6489), 409–412, doi: 10.1126/science.abb3405 (2020).

41. Schäfer, A. et al. Therapeutic treatment with an oral prodrug of the remdesivir parental nucleoside is protective against SARS-CoV-2 pathogenesis in mice. Science Translational Medicine. 14 (643), eabm3410, doi: 10.1126/scitranslmed.abm3410 (2022).

42. Sheahan, T.P. et al. An orally bioavailable broad-spectrum antiviral inhibits SARS-CoV-2 in human airway epithelial cell cultures and multiple coronaviruses in mice. Science Translational Medicine. 12 (541), eabb5883, doi: 10.1126/scitranslmed.abb5883 (2020).

43. Owen, D.R. et al. An oral SARS-CoV-2 Mpro inhibitor clinical candidate for the treatment of COVID-19. *Science (New York*, N.Y.). 374 (6575), 1586–1593, doi: 10.1126/science.abl4784 (2021).

44. Vuong, W. et al. Improved SARS-CoV-2 Mpro inhibitors based on feline antiviral drug GC376: Structural enhancements, increased solubility, and micellar studies. European Journal of Medicinal Chemistry. 222, 113584, doi: 10.1016/j.ejmech.2021.113584 (2021).

45. Vuong, W. et al. Feline coronavirus drug inhibits the main protease of SARS-CoV-2 and blocks virus replication. Nature Communications. 11 (1), 4282, doi: 10.1038/s41467-020-18096-2 (2020).

46. Zuckerman, N.S., Bucris, E., Keidar-Friedman, D., Amsalem, M., Brosh-Nissimov, T. Nirmatrelvir Resistance—de Novo E166V/L50V Mutations in an Immunocompromised Patient Treated With Prolonged Nirmatrelvir/Ritonavir Monotherapy Leading to Clinical and Virological Treatment Failure—a Case Report. Clinical Infectious Diseases. 78 (2), 352–355, doi: 10.1093/cid/ciad494 (2024).

47. Miura, T. et al. In vitro selection of macrocyclic peptide inhibitors containing cyclic γ2,4-amino acids targeting the SARS-CoV-2 main protease. Nature Chemistry. 15 (7), 998–1005, doi: 10.1038/s41557-023-01205-1 (2023).

